# Biological Self-organisation and Markov blankets

**DOI:** 10.1101/227181

**Authors:** Ensor Rafael Palacios, Adeel Razi, Thomas Parr, Michael Kirchhoff, Karl Friston

## Abstract

Biological self-organisation is a process of spontaneous pattern formation; namely the emergence of coherent and stable systemic configurations that distinguish themselves from their environment. This process can occur at various spatial scales: from the microscopic (giving rise to cells) to the macroscopic (the emergence of organisms). Self-organisation at each level is essential to account for the hierarchical organisation of living organisms (organelles within cells, within tissues, within organs, etc.). In this paper, we pursue the idea that Markov blankets – statistical boundaries separating states that are external to a system from its internal states – emerge at every possible level of the description of the (living) system. Through simulations, we show that the concept of a Markov blanket is fundamental in defining biological systems and underwrites the nature and form of interactions between successive levels of hierarchical structure. We demonstrate the validity of our argument using simulations, based on the normative principle of variational free energy minimisation. Specifically, we adopt a top-down approach to provide a proof of concept for the claim that the self-organisation of Markov blankets (and blankets of blankets) underwrites the self-evidencing, autopoietic behaviour of living systems.

## 1. Introduction

This paper is about the essential role played by Markov blankets in (self-organised) living systems. A Markov blanket is a statistical boundary that separates two sets of states (e.g. a cellular membrane separating intracellular and extracellular dynamics). The Markov blanket precludes direct interactions between internal and external states – any interactions are mediated through the states that constitute the Markov blanket. As we shall see below, this separation is a fundamental property of living systems because their very existence implies the presence of a boundary that distinguishes inside (i.e. self) from the outside (i.e. environment). Living systems maintain the integrity of these boundaries, in the face of an everchanging environment. This means that life has evolved mechanisms for the generation, maintenance, and repair of Markov blankets.

A system endowed with such mechanisms connotes an autopoietic organisation; it is capable of autonomously producing its own components, in particular its boundaries, [1], [2]. This autonomy does not imply isolation from the environment, which – on a thermodynamic account – is needed to provide a constant energy supply [3]. Therefore, living organisms are operationally closed, while presenting as thermodynamically open [4]. The interaction between system and environment is then mediated by the boundary. Notably, this coupling is non-trivial; in the sense that the organism must actively realise an ‘informational control’ of the environment (i.e., possess a teleology), by filtering, canalising and categorising signals that carry information about their external causes [4]. At the same time, the (statistical) boundaries must contain the machinery that allows the system to act on the external world; namely, active states. In short, definitive borders are essential for living systems, as any dynamics that happens within and between systems can only take place in virtue of their existence [5].

Living organisms are complex systems, denoted by non-linear interactions between multiple hierarchically arranged and nested components [6], [7]. As such, characterising how they self-organise requires not only an understanding of how single components couple to each other, but also of how microscopic and macroscopic components interact. This requires us to acknowledge the existence of top-down influences on the low level dynamics [8] and *vice versa*.

Self-organisation has been addressed extensively in theoretical biology using tools from statistical thermodynamics and information theory to explain how biological systems resist a natural tendency to disorder. This holdout is an apparent violation of the second law of thermodynamics, or, more precisely, the fluctuation theorems for non-equilibrium systems [4], [9]–[12], which states that the probability of their entropy decreasing itself decreases exponentially with time (and scale). A prominent line of work within this framework sees living organisms as constantly minimising an upper (free energy) bound on their self-information (i.e., negative log likelihood of sensed states). This imperative is motivated by the fact that biological systems have to maintain sensory states within physiological bounds. This means the Shannon entropy (dispersion) of sensory states is necessarily bounded [13]. In this setting, the Shannon entropy is the path or time average of self-information; also known as *surprisal* or *surprise*. In short, self-organisation can be regarded as synonymous with systems that place an upper bound on their self-information or surprise. In current (variational) formulations of self-organisation – that emphasise its sentient or inferential aspect – living organisms are understood as placing a (free energy) bound on surprise, rather than reducing surprise directly.

These arguments rest upon mild ergodicity assumptions (implicit in the fact that the sorts of systems we are interested in have characteristic measures that persist over time). Ergodicity implies that, over a sufficiently long period, the time spent in a particular location of state-space is equal to the probability that the system will be found at that location when sampled at random [5]. If this probability measure is finite, it means that any system will revisit certain states (or their neighbourhoods) time and time again. It is this peculiar and special behaviour that underlies self-organisation; namely, the existence of an attracting set of states that endow living systems with characteristic behaviours that occur repeatedly.

The existence of an attracting set means that one can interpret the long-term average of surprise of sensory states as the average surprise conditioned on the system over all possible sensations, which is equal to their entropy. This means that minimising the bound on surprise minimises entropy, or the dispersion of sensory states [14]. Crucially, because surprise is (negative) Bayesian model evidence, minimising free energy is the same as maximising a lower bound on the evidence for an implicit model of the causes of sensations. In other words, the system can be regarded as a model of its environment [15], and will try to gather evidence for its own existence. This has been called self-evidencing [16]. It follows that – by minimising free energy – biological systems place an upper bound to the entropy of their sensations, while inferring their causes; this is also known as *active inference* [17], and is closely related to other formulations of the perception-action cycle in other disciplines, like embodied cognition [18], artificial intelligence [19], and cognitive neuroscience [20]. In short, *self*-organisation entails the bounding of *self*-information that can be cast as *self*-evidencing.

If a biological system did not minimise (a free energy bound) on surprise it would cease to exist, as the entropy of its sensory states would increase indefinitely. In other words, it would dissipate, decay, dissolve or die. Friston [5], demonstrated that (almost) any (ergodic random dynamical) system endowed with boundaries (Markov blankets) is autopoietic (self-organising). In other words, the system appears to minimise free energy and engages in active inference and thereby actively maintain its functional and structural integrity. Both a heuristic proof and proof of principle were provided to support this claim. The latter comprised a simulation of a primordial ‘soup’ or ensemble of subsystems; each with its own physical and electrochemical states, coupled through short-range interactions. The equations of motion of the subsystems were integrated until nonequilibrium steady state. This allowed one to identify a Markov blanket separating some internal states from their environment – based on statistical dependencies between subsystems that emerged during the evolution. This work effectively used a bottom-up approach to show that self-organisation entails the emergence of Markov blankets that can be cast in terms of active inference or self evidencing.

Here, we provide a proof of concept that complements the work described above. In contrast to the bottom-up approach, we adopt a top-down view; building upon the free energy formulation of pattern formation [21]. This means that we start with subsystems whose dynamics possess a Markov blanket as an attractor. We then integrate the system until it self-organises into a stable configuration. We then consider hierarchical systems; namely, configurations of configurations (i.e., blankets of blankets) that could, in principle, be extended indefinitely. We argue that, given local interactions, Markov blankets are an essential feature of any biological system. More specifically, we test the following hypothesis: if the maintenance of Markov blankets – that underwrite existential form – can be cast as self-evidencing, then self-organisation should be an emergent property of subsystems that ‘believe’ they participate in – or are enclosed by – a Markov blanket. Because Markov blankets are defined by conditional independence; the requisite ‘beliefs’ can be specified simply in terms of communication or signalling between subsystems. In other words, it should be possible to prescribe hierarchical self-organisation purely in terms of whether or not any element of an ensemble can influence – will be influenced by – another element, depending upon their role as a Markov blanket or internal state at the next hierarchical level.

In such systems, self-organisation should, in principle, lead to the formation of nested (statistical) boundaries as we ascend the hierarchy. Here, we associate random dynamical systems with living organisms. Of course, this is a tremendous simplification, motivated by the fact that the systems under consideration are complex (i.e., non-linear and hierarchical) and organised independently of any apparent external gradient: in other words, pattern generation starts as soon as the system exists.

This paper is organised as follows: in section 2 we introduce the concept of Markov blankets and argue that any biological system has to conform to such an organisation. In section 3, we follow the evolution of a random dynamical system – endowed with a Markov blanket – via the principle of free energy minimisation (i.e. self-evidencing through active inference). This illustrates the autopoietic nature of systems that, through the dynamics of their internal and active states, resist a natural tendency to disorder. In sections 4 and 5, we describe simulations of self-organisation at two hierarchical levels; these furnish a proof of concept for self-organisation of Markov blankets of Markov blankets. We conclude with a discussion of future considerations in section 6.

## 2. Markov blankets

The notion of Markov blankets was originally proposed in the context of Bayesian networks or graphs [22], where it refers to the parents of the set of states (that influence it), its children (that are influenced by it), and the children’s parents. The Markov blanket defines the conditional independencies between a set of states (the system) and a second set of states (the environment). This concept can be gracefully translated into a biological setting: for example, the internal milieu of a cell represents the internal states, the environment external states, and the plasmalemma is the Markov blanket through which communication between intracellular and extracellular states is mediated [5], [23]. Crucially, the Markov blanket can be decomposed in sensory and active states, which are and are not children of the external states, respectively. Thus, the existence of a Markov blanket *S* × *A* induces a partition of states in *x* ∈ *X* = *Ψ* × *S* × *A* × Λ; external states act on sensory states, which influence, but are not influenced by internal states. Internal states couple back through active states, which influence but are not influenced by external states (Table 1). This circular causality is clearly reminiscent of the perception-action cycle [5].

**Table 1.**
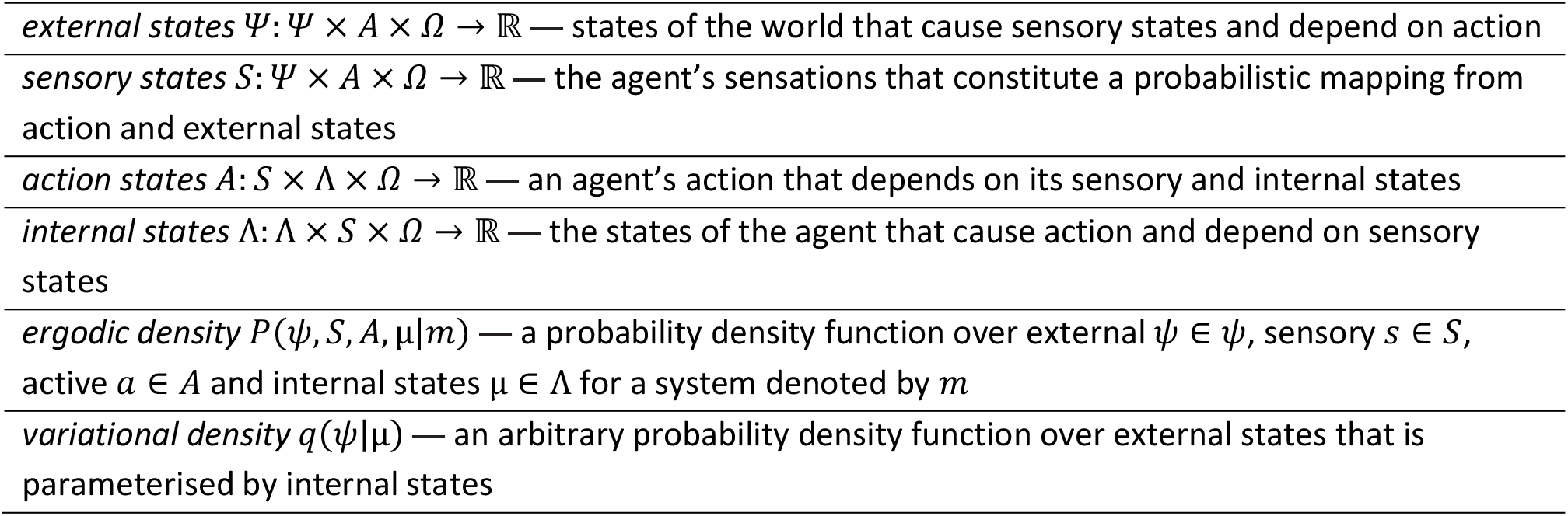
Definition of the tuple (*Ω, Ψ, S, A*, Λ, *p, q*) underlying active inference. a sample space *Ω* or non-empty set from which random fluctuations or outcomes *ω* ∈ *Ω* are drawn

Why is the presence of a Markov blanket – and the resulting partition of states in four sets – so important? To understand this, let us consider a system, composed of different components; where long-range (e.g. electromagnetic) interactions are possible. Each state will interact with all others, irrespective of its spatial position. In this system, every component will eventually become indistinguishable from the others, because the fully interconnected nature of the system precludes any statistical separation of one component from another (Figure 1a). In order to engender statistical structure, coupling has to be limited. This is possible by introducing short-range interactions, whereby coupling becomes spatially dependent (Figure 1b). However, in such a system, the existence of two distinct sets of states is only possible if they are far apart, so that interactions are precluded.

**Figure 1.**
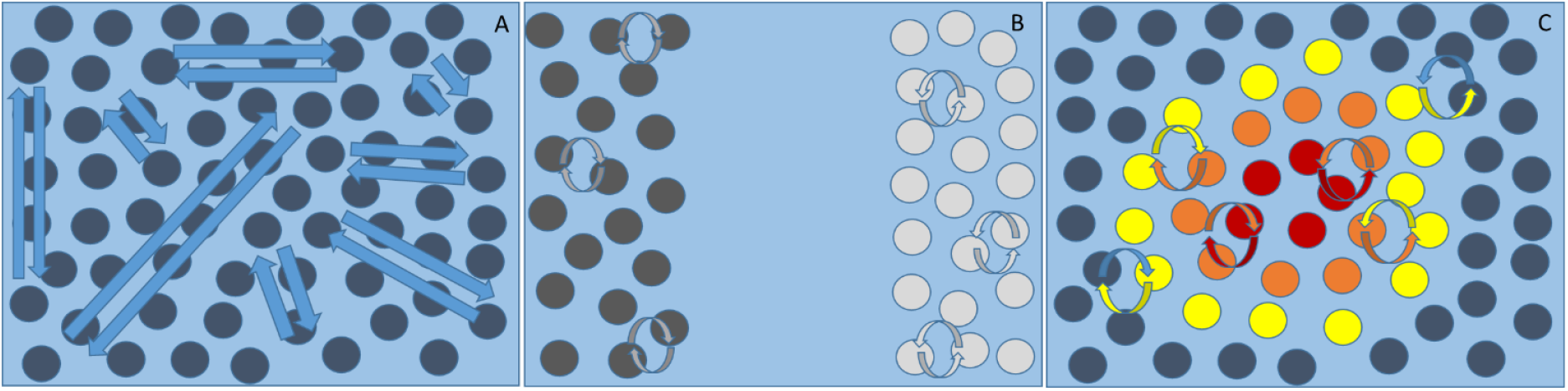
*System comprising interacting components. In (a) spatially-independent coupling among subsystems is mediated by long-range interactions. In the first (left) panel all states influence each other, and are therefore indistinguishable. In (b) only short-range interactions are allowed; thus coupling among subsystems is spatially dependent. However, two sets of states exist only because of spatial separation: they are effectively independent. In (c), internal (red) and external (blue) states can be distinguished in virtue of the separation operated by a third set; namely, the Markov blanket, composed of sensory (yellow) and active (orange) states. External states can influence internal states only by acting on sensory states. On the other hand, internal states couple back to external states through active states*.

However, in an interesting system, segregation (i.e., self-organisation) persists in the presence of communication. In other words, a system segregates from the environment, but remains (statistically or energetically) coupled to it. Ultimately, we arrive at a third case (Figure 1c). In this case, two sets of states exist not just because of their spatial separation, but in virtue of the existence of a third set, namely the Markov blanket. These blanket states comprise sensory and active states, mediating the vicarious coupling between internal and external states. States of the Markov blanket surround one set of (internal) states, and isolate it from the second set of (external) states. Now, external states can influence internal states only through sensory states. At the same time, internal states couple back through active states. In short, the Markov blanket provides a statistical insulation whereby *internal* states can be regarded as *insular* states. This concludes our description of the minimal conditions necessary for a system as simple as a bipartite universe to exist.

Now we take a step back and consider the ensemble of internal states and their Markov blanket as a unitary (multidimensional) state. In order to form some sort of meaningful separation at this macroscopic level of description, a new, bigger Markov blanket has to emerge, whose sensory and active states – and the internal states insulated within – will each be composed of a smaller Markov blankets (and internal states). Hence, the formation of Markov blankets at any level of the hierarchical organisation (that underwrites the structure of biological systems) is intimately linked to the maintenance of Markov blankets ‘all the way down’ (Figure 2).

**Figure 2.**
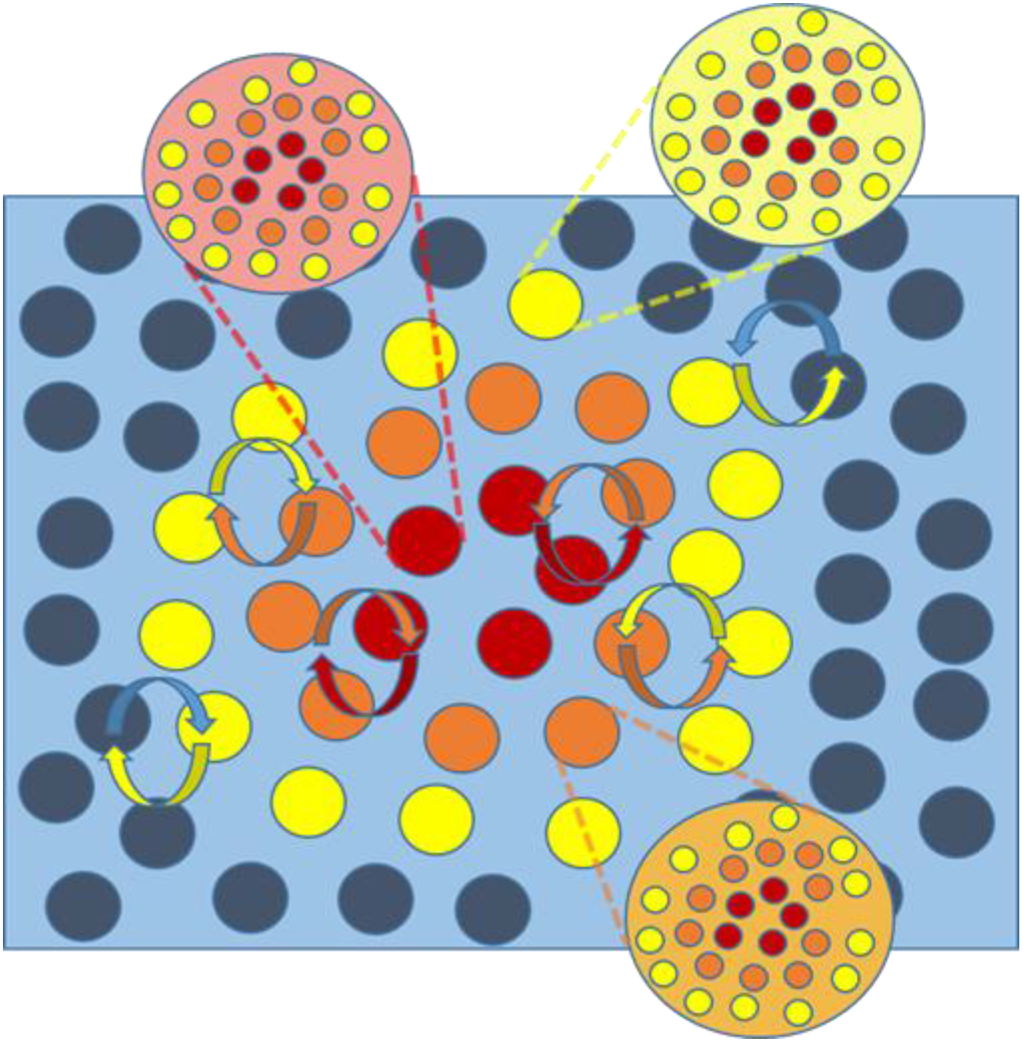
*Markov blanket of Markov blankets. We now broaden the perspective, and consider each Markov blanket (and internal states) as a collective state. Again, given short-range interactions, the only way for a system to exist at this new level is to be separated from its environment by a Markov blanket. The hierarchical nature of this system induces Markov blankets of Markov blankets; the emergence of Markov blankets occurs hierarchically: the big Markov blanket (and its internal states) is constituted by smaller Markov blankets (and their internal states)*.

On this view, it is clear that the self-organisation of living organisms has to feature the emergence of boundaries that define an internal space, separating it from the environment, while keeping them indirectly coupled. Furthermore, this self-organisation is a recursive process that spans all levels of hierarchical organisation. In what follows, we provide a proof of concept for this argument by simulating the hierarchical self-organisation of Markov blankets.

At this point it is interesting to note that disabling long-range interactions and retaining only local coupling simplifies the fully interconnected picture of Markov blankets mediating the perception-action cycle above: sensory and active states do not act directly on internal and external states, respectively. Nonetheless, internal and external states remain insulated from one another through the Markov blanket; thus, the conditional independences essential for the existence of the system (i.e., its Markov blanket) are preserved.

## 3. Dynamical systems, self-organisation and self-evidencing

We will be dealing with random dynamical systems, whose state equations are described by random differential equations of the following form:

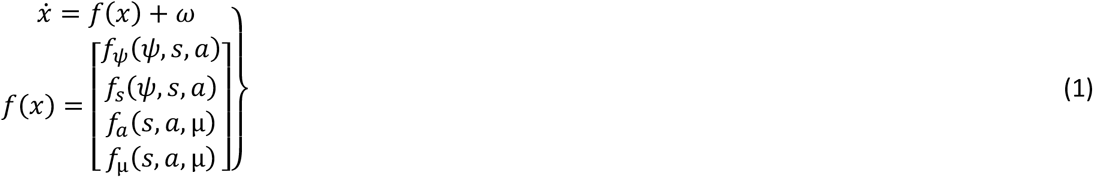

These equations can be regarded as describing the evolution of states of a system and its local environment, in terms of the motion of states *f*(*x*), subject to random fluctuations *ω*. The distinction among external, sensory, active and internal states is formalised in the second equation by the dependencies implied by the Markov blanket. External states can only be accessed by internal states through the Markov blanket, and are therefore usually called hidden (or latent) states. These states can be interpreted as the ‘true’ states of the embodied system, comprising both external conditions (i.e. the environment) and physiological conditions (e.g. body temperature or pressure). In both cases, these (external) states can be seen by internal states (e.g., a brain or intracellular organelle) only through the Markov blanket.

Following the formulation of [24], we use the Helmholtz decomposition, such that we can express the flow of states in terms of a divergence-free component and a curl-free descent on a geometrical space determined by a scalar *Lagrangian L*(*x*) that corresponds to the self-information or surprise associated with any state.

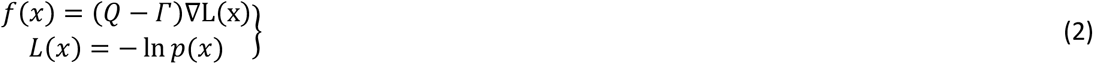

The diffusion tensor Γ is half the covariance (amplitude) of the random fluctuations, and *Q* is an antisymmetric matrix that satisfies *Q*(*x*) = –*Q*(*x*)^*T*^. Because the system is ergodic, it will converge toward a set of states, called a pullback or *random global attractor*, whose associated probability density we will call an ergodic density [25], [26]. It is straightforward to show that *p*(*x*|*m*) = exp(–*L*(*x*)) is the ergodic solution of the Fokker-Planck equation [27], also known as the Kolmogorov forward equation [28] describing the density dynamics. This means that we can express the flow in terms of the ergodic density

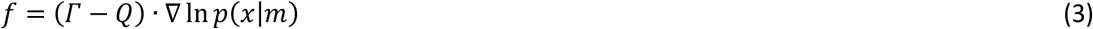

This equation means that the states of an ergodic system effectively perform a gradient ascent on the ergodic density. This in quite revealing because it shows that the system’s flow counters the dispersive effects of random fluctuations – by flowing towards the pullback set of states. This also applies to the motion of internal and active states

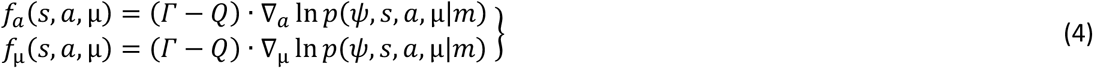

These equations are the homologues of (3) for internal and active states. They say that their flow performs a generalised gradient ascent on the ergodic density that describes the internal states and Markov blanket of any system. In other words, the system is autopoietic, as its characteristic probability density over states is maintained by the motion of its own internal and active states.

The flow of the states therefore describes a gradient ascent on the ergodic density. Analogously, in the setting of the stochastic thermodynamics of non-equilibrium steady states, the system is minimising its free energy [29]. Although the ergodic density exists, it cannot be computed explicitly by the system, because this would require access to external states that are hidden to the internal states. However, it is possible to use an alternative formulation that allows a description of the flow in terms of a gradient descent based on the variational free energy associated with a generative model of the system in question [21]:

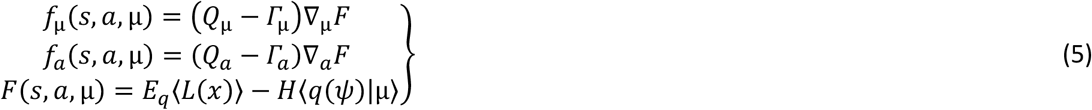

Here, the flow of internal and active states constitutes a gradient on variational free energy, which is a function of states that are available to the system. This follows because free energy depends on a variational density *q*(*ψ*|μ)over external states that is parameterised by internal states, and a generative model *p*(*ψ, s, a*, μ|*m*), which is the system itself [21].

Under this formulation of density dynamics, internal states will appear to *infer* external states. The third equality expresses free energy as the self-information (i.e., negative log evidence for the model) expected under the variational density minus the entropy of the variational density. This means that internal and active states maximise the joint probability density – expected under the variational density – over states conditioned on the system or model in question. Moreover, internal states will also reduce free energy by parameterising a variational density over external states with maximum entropy, in accordance with Jaynes’ principle of maximum entropy [30]. Although not our focus here, the variational density becomes the posterior distribution of hidden or external states, given blanket states, when variational free energy is minimised. In this sense, the internal states encode posterior ‘beliefs’ about external states; despite never seeing them directly.

The free energy formulation of a system’s dynamics allows us to prescribe the ergodic density in terms of a generative model. In other words, we could write down some equations of motion and interpret the resulting ergodic density as the surprise associated with an unknown generative model (the top-down approach). Alternatively, we can write down a generative model and derive the dynamics according to Equation (5) as a gradient descent on the free energy equivalent of surprise. In what follows, we will simulate self-organisation by specifying a model about the causes of the system’s sensory states – and by specifying the environmental dynamics generating those sensations.

This means we need to write down the generative model *p*(*ψ, s, a*, μ|*m*) of the system in terms of the dynamics *f_ψ_*(*ψ,s,a*) and *f_s_*(*ψ,s, a*) of the environment and how sensory states are generated. Interestingly, the generative process and model do not have to be isomorphic: the generative model has only to approximate the generative process to minimise free energy. The generative model is usually expressed in terms of random differential equations and nonlinear functions with a hierarchical form (in this paper, we will only need to specify nonlinear functions):

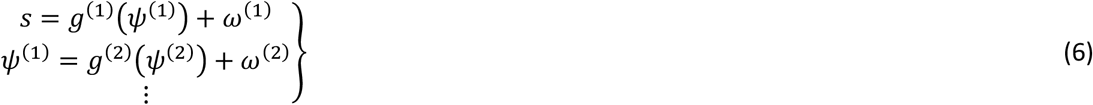

Under Gaussian assumptions about random fluctuations ω, these nonlinear functions prescribe the likelihood and priors over external states, from which the Lagrangian is recovered

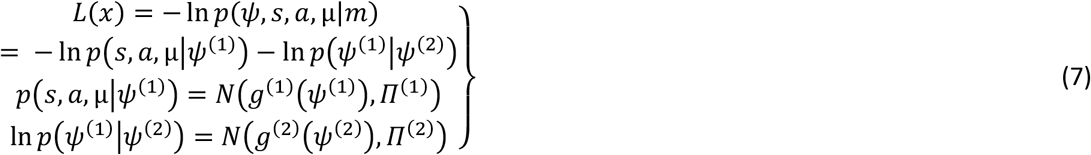

Here, *Π*^(*i*)^ corresponds to the precision or inverse variance of the random fluctuations. In what follows we integrate Equation (5) using the Matlab routine **spm_ADEM.m** in the SPM open source academic software. This generalised filtering or integration scheme uses the Laplace assumption to specify the (Gaussian) form of the variational density, and can be regarded as a generalised Bayesian filter. This follows because the variational density *q*(*ψ*|μ) over external states approximates the posterior density *p*(*ψ*|*s, a*, μ), as noted above. See [24] and [13] for details.

In summary, we will use a standard (generalised) Bayesian filtering scheme to simulate self-organisation within a random dynamical system. Using a Bayesian filtering scheme means that we get the requisite partition into external, sensory, internal and active states for free. Furthermore, we can specify the form of the ergodic or nonequilibrium steady-state density in terms of a Lagrangian – by formulating the flow of internal and active states in terms of variational free energy – that can be specified in terms of a generative model. The question now is: what sort of model leads to hierarchical self-organisation?

## 4. Self-organisation of an ensemble

In what follows, we present two sets of simulations. The first considers the self-organisation of ensemble of cells, where each cell possesses its own Markov blanket. The second simulations consider ensembles of ensembles to illustrate hierarchical self-organisation; namely, the self-assembly of Markov blankets of Markov blankets of Markov blankets. Crucial to these simulations is the use of simple generative models, embodying the prior ‘belief’ that each member plays the role of an internal, active or sensory state within the ensemble. In other words, Markov blankets at one level of organisation possess prior ‘beliefs’ there is a Markov blanket partition at the level above. This is easy to specify because each role just depends upon the influences each member of the ensemble can or cannot exert on the others. Furthermore, the only hidden state each member needs to infer is which role it plays at the higher level. We will see that this minimal set of prior beliefs (and subsequent self-evidencing) results in the formation of Markov blankets within the ensemble. The ensuing self-similar organisation can, in principle, be extended to any number of hierarchical levels. We will illustrate this below using 16 cells, each with their own Markov blanket, that organise into a cellular group or assembly, with its own Markov blanket. We then consider an ensemble of ensembles that organises itself into a little organ encompassed in another Markov blanket.

The first simulation illustrates the self-organisation of an ensemble or multi-agent system. Each component (e.g. cell) interacts with other cells; in a process that eventually leads to a stable configuration with a boundary separating internal cells from their external milieu. This simulation draws on previous work that simulated morphogenesis [21]. In this setting, self-organisation was simulated by minimising the variational free energy of each cell until they attained a prescribed morphology. This morphology was achieved through spatially dependent (e.g. chemical) signalling – so that every cell sensed every other cell in a way that was consistent with their generative models. The morphology was inscribed in beliefs common to all cells, about cell identity, sensation and secretion. Each cell was interpreted as a Markov blanket surrounding internal states: the action (active states) of a cell was the cause (i.e., external states) of the sensations (i.e., sensory states) of the remaining cells. At the beginning of pattern-formation, cells were undifferentiated, because they were uncertain about their identity in the target morphology. As selforganisation unfolded, each sub-system inferred a unique identity, location and what they should sense at that location. When every cell was in the right place, these inferences were fulfilled; thereby minimising the free energy (i.e., self information or surprise) of every cell.

In more detail, this inference – in analogy to intracellular cascade signalling and epigenetic mechanisms – was driven by the minimisation of free energy. By generating identity-dependent predictions (e.g. genetic and epigenetic expression) about sensations, every cell moved around and generated extracellular signals until its predictions were confirmed. Predictions about sensations caused by the other (e.g. extracellular signalling) and its own action (e.g. secretion and position) were dictated by prior beliefs (in the generative model) about the role of each cell in the target morphology. These prior beliefs were the same for every cell (c.f., pluripotential or stem cells). In other words, based on its identity, each cell had particular expectations about its sensory states. Because sensations were caused by other cells, surprise could only be minimised when every member of the ensemble had inferred a unique role within the ensemble. In short, priors established a point attractor for the ensemble dynamics, in terms of a free energy minimum.

In the present work, we use the same strategy: we simulate self-organisation of a multi-agent system, whose components – coupled through spatially decaying (e.g. chemical) signals – minimise variational free energy, based on a generative model describing how causes generate sensations. Again, as the external states of each component are the active states of other cells, the system organises in a pattern that enables each cell to predict signals from its companions as precisely as possible. However, in the current simulations, the prior beliefs were much simpler: they specify signalling and three possible types of cell, so that each cell only had to infer what type of cell it was. This means that there are no target positions or target morphology. The only prior constraints are beliefs about the intracellular and extracellular sensations for the three cell types. Crucially, these priors conform to the conditional dependencies and independencies entailed by a Markov blanket: active cells can sense and be sensed by both sensory and internal states, whereas sensory and internal states that do not interact.

From the perspective of the ensemble as a whole there are no external states, which would interact only with sensory states. This is an important point, because self-organisation is auto-referential here – as it does not require coupling with an external environment. The ensuing self organisation leads to a spatial pattern, wherein components of the system are organised in a predictable fashion. Such a pattern is inscribed in the (e.g., genetically encoded) expectations about sensations of the components of the system. More precisely, priors specify the form of the generative model, which corresponds to the free energy landscape, thus defining the sets of attracting states towards which the dynamics of the single components converge [21]. In short, priors of a model dictate how the system self-organises, granted that such a model leads to prediction error minimisation.

We now describe our simulation setup. The system comprised sixteen cells, which can become one of three types of cells that play the role of states at the next hierarchical level; namely, internal, active and sensory states. Each cell type (believes it will) secrete a unique chemical signal and communicate according to the conditional independencies required by a Markov blanket (see Table 2). The external states of each cell comprised its location 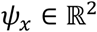 and the chemical signals 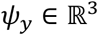 released. This can be expressed as

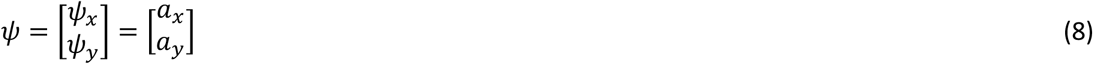

**Table 2.**
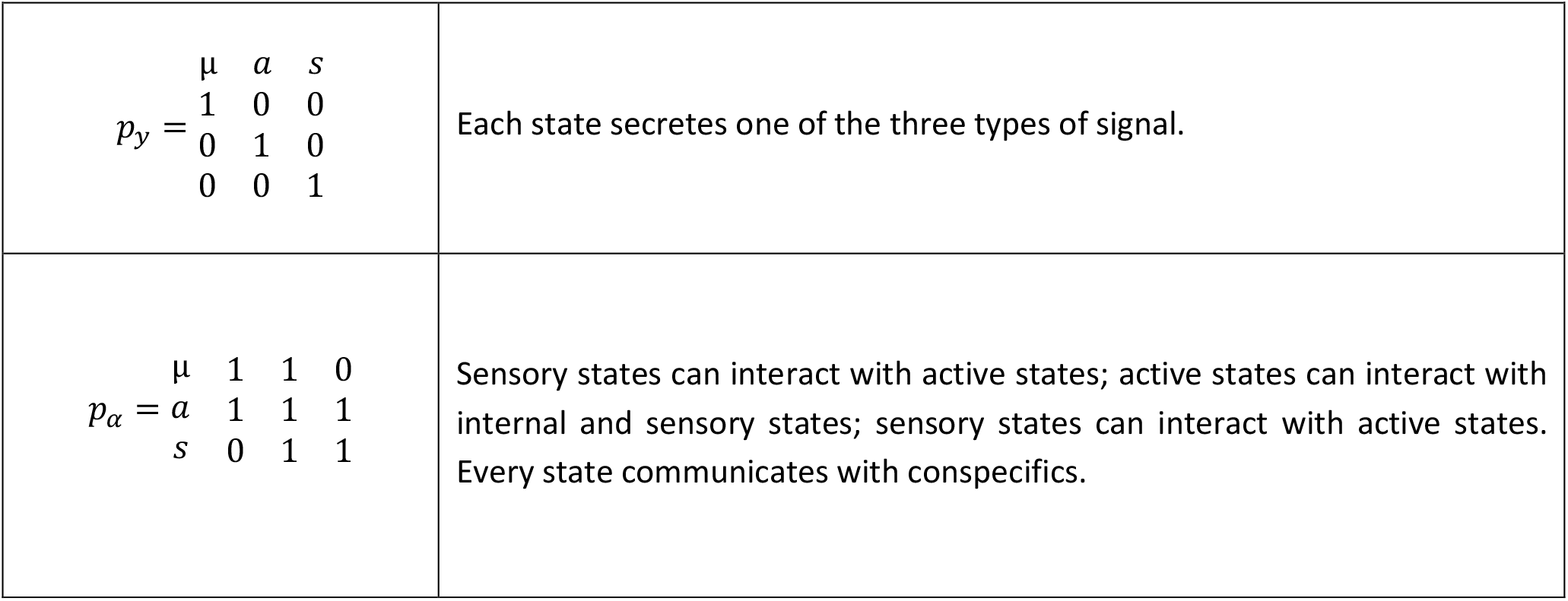
Prior beliefs characterising Markovian dependencies and independencies

For simplicity, the location and signalling were also taken to be the active states of any given cell. Its sensory states are the sensed intracellular (produced by itself) and extracellular (produced by other cells) signals. The latter is a function of distance, as we assume concentrations of secreted chemicals decreased exponentially over space. This can be expressed as

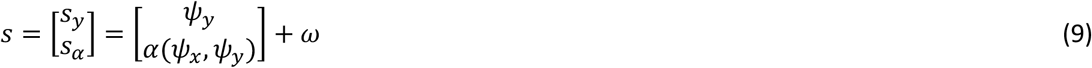

Here, the sensory noise *ɷ* had a high precision of exp(16). The sensed extracellular signals are returned by the function *α*(*ψ_x_, ψ_y_*), which accounts for the spatial decay of chemicals; where the extracellular sensations of the *i*^th^ cell are given by

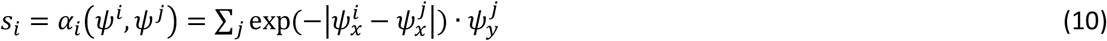

Here, *j* indexes all cells other than the *i*^th^ agent. Each cell generates predictions based on the same generative model, which specifies the mapping from hidden states – namely, the *type* of the cell *ψ_i_* – to sensations. The type is then the only hidden state that the cells have to infer, which is parameterised by their internal states μ_*i*_. Based on beliefs about its type, each cell then generates predictions about intracellular and extracellular sensations:

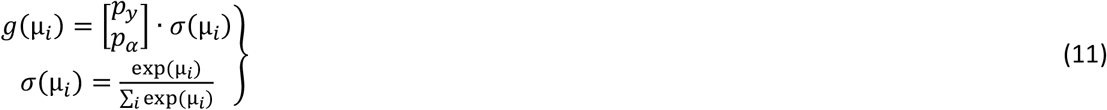

Here, *p_α_* and *p_y_* are prior beliefs about secretion and sensation given the type of cell (see Table 2), while *σ*(μ_*i*_) is a softmax function that returns the expectations about the cells type. The resulting dynamics of internal and active states of each cell (suppressing higher order terms for clarity) can be expressed as follows:

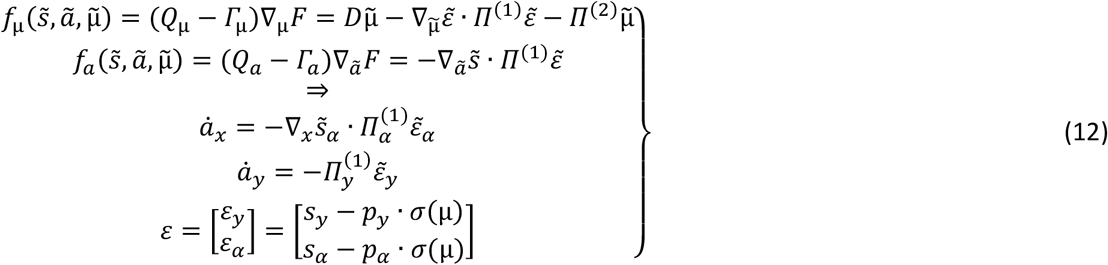

Here *ε* = *s* – *g*(μ) is called a prediction error, and *Π*^(2)^ is the precision (with a log precision of minus four) of a Gaussian prior over internal states that parameterise posterior ‘beliefs’ about external states. The ~ notation denotes generalised motion: see [3]. Equation 12 shows that internal and active states minimise (variational) free energy. Under the Laplace assumption, this effectively reduces to prediction error minimisation. Thus, internal and active states perform a descent on prediction error gradients [14]. Under these equations of motion, cells infer their identity based on sensations, while secreting according to their role as the ensemble evolves. At the same time, cells move to reach a position where extracellular inputs can be best predicted.

The results of an exemplar simulation are summarised in Figure 3. Self-organisation leads the ensemble to assume another cell-like morphology with internal cells in the middle, encircled by active cells, surrounded in turn by sensory cells. Because there are no prior beliefs either about the location or about the number of cells per type, this pattern constitutes an emergent property. This is because the prior beliefs define (statistical) coupling among members of the ensemble, while leaving its topology unspecified. In other words, the final morphology of the ensemble is an emergent property of the spatially dependent interactions among agents and conditional independencies consistent with a Markov blanket. Crucially, the ensuing self-organisation produces a spatial structure that resembles the most elementary biological unity – a cell. In short, the cell-like organisation of the ensemble emerges from intracellular signalling in a way that does not require any morphological priors. These results therefore support the idea that (e.g., genetic) beliefs entailed by a Markov blanket are sufficient for the emergence of structures with statistical boundaries that distinguish internal states from the external milieu. In turn, this suggests that priors that embody Markovian dependencies may play an essential role in the self-organisation of biological systems.

**Figure 3.**
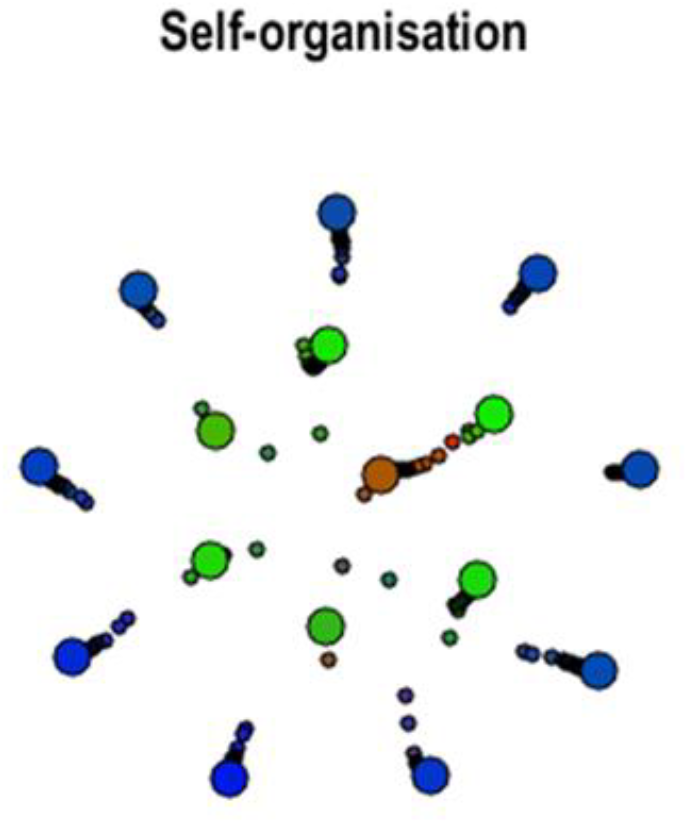
*Self-organisation at a particular level. This figure illustrates the (final stage of) selforganisation of an ensemble comprising sixteen ‘cells’, whose internal and active equations of motion describe a gradient descent on prediction error, relative to sensory states expected by each member of the ensemble. Every member is endowed with the same prior beliefs about what they should signal and sense, depending upon their type (which has to be inferred on the basis of what they sense). These priors ultimately prescribe a point attractor for the dynamics of the ensemble. Each cell can then assume one identity or type and behave accordingly, while moving to a location that fulfils its predictions about its extracellular signals. The emergent morphology of the ensemble has a cell of cells form, with an internal (red) cell in the centre, surrounded by a membrane of active (green) cells in the middle, and sensory (blue) cells on the periphery. This is the spatial pattern that best fulfils the prior beliefs of all the constituent cells*.

The simulation presented above illustrates the biological importance of Markov blankets in a simple but plausible world where only local interactions are permitted, in which prior beliefs (e.g., a genetic code) have learned that, in order to exist, a living system has to self-generate boundaries that separate it from – and mediate the coupling with – its environment. As in real biological systems, the constituents interact with each other, leading to signal cascades. The (e.g., epigenetic) signalling rests on inference about the type or role each cell should play, where action (e.g., chemotactic signalling) realises that role. Cells then differentiate, based upon their prior beliefs (e.g., genetic code). In essence, the ensemble reaches a steady state characterised by an internal milieu, which exists – in virtue of assembling its own Markov blanket – as integral part of the system. One might imagine that genes specify Markovian affordances to produce hierarchical structures; such as organs, tissues, organisms and so on. On this view, self-organisation is then a recursive process that engenders, at every level, the emergence of Markov blankets.

## 5. Self-organisation: ensemble of ensembles

In the final set of simulations, we simulated hierarchical self-organisation in sixteen ensembles composed of sixteen cells each. To investigate the autonomous organisation of (256) cells at two levels, every cell is equipped with the same priors about their local and global identity, that is, they share beliefs about possible roles at both ensemble (local) and ensemble of ensemble (global) level. Cells then can infer their identity both at the local and global level, simultaneously. Put simply, each cell now had two sets of hidden states – and prior beliefs – pertaining to their role at the local and global level. Crucially, these priors are identical, and are the same as used in the previous simulation; namely, they prescribe conditional independencies that are mandated by a Markov blanket at each level in the self-similar fashion:

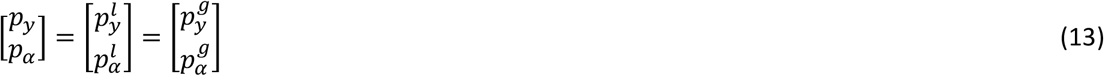

Here, the superscripts denote the local (ensemble) and global (ensemble of ensemble) level. The only additional piece of information required in this simulation is how the two levels couple to each other. For simplicity and computational expediency, we model the microscopic dynamics (cells within an ensemble) of only one ensemble of sixteen cells, whereas for the remaining (fifteen) ensembles, we assume that the average behaviour conforms to the local dynamics of the simulated ensemble. This is a mean field approximation in the sense that we discount local fluctuations within each ensemble and assume their average behaviour is ‘seen’ by any single ensemble. This allows us to simulate the coupling of sixteen cells of the fully simulated ensemble with other fifteen ensemble means (without simulating the other 15 ensembles explicitly). In summary, this simulation illustrates how sixteen cells self-organise in an ensemble that in turn self-organises with other fifteen identical ensembles, while describing the coupling between the local and global level.

In particular, for the fully simulated *k^th^* ensemble, the global to local extracellular coupling means that it only senses the (simulated) average of all other global signals, while the local to global coupling means that the average over its active states informs the dynamics of the remaining ensembles. Technically, this means:

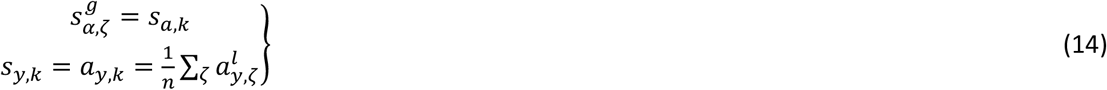

where *ζ* = 1:*n*. The first and second equalities in (14) refer to the extracellular sensing of cells and the intracellular sensation of the ensemble, respectively. In terms of local to global coupling, as they are part of the same ensemble, these predictions will be congruent with each other and cells will therefore act in concert at the global level:

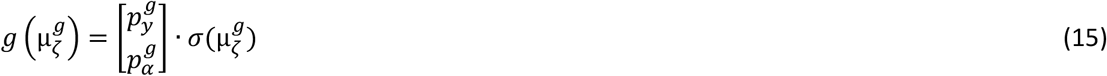

where 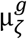 is the expectation about global type for every cell in the ensemble. In summary, sixteen cells locally self-organise in an ensemble, guided by the local priors, while interacting with the remaining fifteen ensembles. This induces a hierarchical self-organisation and pattern formation of Markov blankets within Markov blankets (see Figure 4). Again there are no external states for the ensemble as self-organisation is autonomous – and leads to the emergence of a pattern where the behaviour of each component conforms to the expectations of the others. The lower panels of Figure 4 show the evolution of subtype expectations (i.e., differentiation) at the local (left), and global (middle) level. The lower right panel shows the expectations of a single ensemble (the sixth) about its role at the global level. Here, the sixth ensemble is an active state at the global level. Note the differentiation on both a local and global level; while local expectations about the cells’ role at the global level converge to the same type.

**Figure 2.**
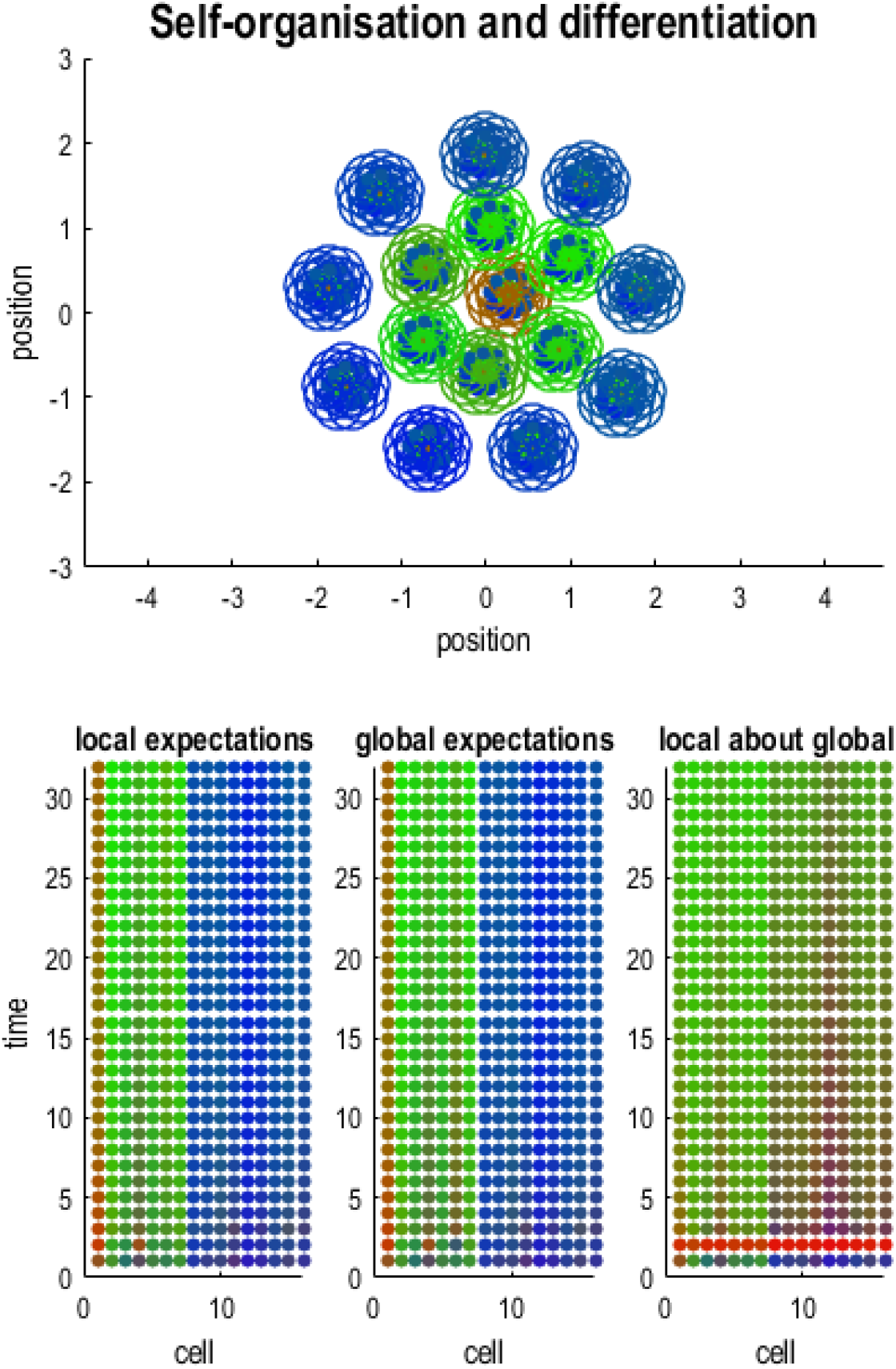
*This figure shows the (final) results of self-organisation of an ensemble of cells, where each constituent of the ensemble is itself a local ensemble. In this example, there are 16 cells at both the global (higher) and local (lower) level. The upper panel shows the final disposition of the ensemble (of ensembles) in terms of the location of cells, and their differentiation (shown in colour: internal – red, active – green and sensory – blue). Note that there are no external states because the external states comprise the Markov blankets of other ensembles. Here, each cell is coded with two colours. The central colour corresponds to expectations about the type of cell in question at the local level, while the peripheral circle encodes expectations at the global level. The key thing to observe here is the emergence of a Markov blanket at both levels. This reflects a particular independency structure, where internal cells do not influence sensory (i.e. surface) cells, in virtue of their separation by active cells. This separation induces conditional independence, because of the limited range of intracellular signals (that fall off with a Gaussian function of Euclidean distance). The lower panels show the same results in a simpler format; namely, the evolution of subtype expectations (i.e., differentiation) at the local (left), and global (middle) level. The lower right panel shows the expectations of a single ensemble (the sixth) about its role at the global level. Here, the sixth ensemble is an active state. Note the differentiation on both a local and global level; while local expectations about the cells’ role at the global level converge to the same type. In these simulations, we used a time step of two units (of arbitrary time) and a second order variational filtering scheme (heuristically, this is a second order generalisation of extended Kalman filtering) with hidden states corresponding to unknown identity in terms of cell type at the local and global level. Please main text for details*.

This simulation exemplifies the absence of a privileged point of view when describing hierarchical selforganisation. The dynamics at every level plays the role of macroscopic states at the level below, and the role of microscopic states at the level above. In self-organisation, interactions among microscopic states inevitably give rise to the macroscopic states that appear to impose constraints on local dynamics. This is formalised in synergetics, and in particular by the slaving principle, which deals with self-organisation and pattern formation in the context of open systems far from thermodynamic equilibrium [31]. In such systems, the fast (stable) dynamics of the microscopic patterns dissipate rapidly as a function of order parameters, where the order parameters are a measure of the macroscopic states that emerge. The basic phenomenology is that these order parameters enslave the dynamics at the level below, which results in an enormous reduction of degrees of freedom. Notably, the emerging macroscopic patterns may sometimes recapitulate microscopic patterns leading to a fractal organisation. This aspect is nicely exemplified by ensembles of oscillators that are coupled together by their average. This generally produces macroscopic dynamics that gives rise to a new oscillator at a larger spatial and slower time scale, while at the same time, each nested oscillator can be regarded as a macroscopic state enslaving the level below [27]. In the same fashion, Markov blankets – that are constituted by Markov blankets – self-organise in ensembles that themselves form Markov blankets at a higher scale.

In summary, we used the same generative model at both levels to exploit the self-similar hierarchical structure that emerges. However, we could have used different generative models at the global and local levels to simulate the morphogenesis of particular organelles that have a different form from their constituent cellular ensembles. We will pursue this in future work.

## 6. Discussion

In this paper, we have considered a variational treatment of self-organisation. Given local interactions, carefully crafted prior beliefs about conditional dependencies and independencies endow a system with a point attractor comprising internal states and their Markov blanket. Moreover, applying the same priors at any hierarchical level leads to the emergence of Markov blankets within bigger Markov blankets. A key feature of the simulations – used in this paper – is the absence of any explicit target morphology within the prior (e.g., genetic) beliefs of the system’s constituents. This is indeed an emergent property, which nonetheless has an apparent top-down causal effect on the blankets below.

Such emergence provides an alternative to the reductionist view established in developmental biology. For example, morphogenesis and development are widely supposed to be primarily guided by local interactions among system’s components, and in particular by morphogen gradients that control gene expression; for example, those established by the Sonic Hedgehog (Shh) protein [32] or by the group of Hox genes [33]. In both cases, concentration gradients underwrite position-dependent differentiation of tissues in the vertebrate’s central nervous system. In addition, Hox genes follow a colinearity rule; in that their respective position in the chromosome parallels the sequence followed by their expression along the anterior-posterior axis in the body. Therefore, not only can genes encode positional information but they can do so explicitly – with a one-to-one correspondence.

On the other hand, our simulations point more towards the idea that a spatial pattern might not be necessarily specified by local interactions and genetic positional information [34]: in the scenario above, target morphology is a consequence of top-down constraints hence the result of interaction among nested levels. This sort of top-down causation, whether real or simplistic, offers nonetheless an efficient way to describe (and manipulate) self-organisation of complex systems.

The second contribution of this work is the extension of self-organising system to potentially higher and higher hierarchical levels. As mentioned above, biological systems are – by definition – hierarchical in their organization [6]. However, this type of complexity is not only a trait pertaining to the living domain; selforganisation (of Markov blankets) can occur at all scales, irrespective of whether it is overtly biological or not [3]. It is indeed possible to imagine systems in which biological organisms occupy only some levels in the hierarchy, but Markov blankets can be found at every level; for example, consider our society and distinct communities within it ??(Kirchhoff et al., submitted)??. Therefore, although the present simulations are strictly anchored in the living realm – because of the genetic flavour of prior beliefs and in the way we have framed them – they speak to an argument that pervades potentially all possible fields of study that concern complex, non-equilibrium (open) systems.

## 7. Conclusion

This work suggests that the Markov blanket is a fundamental characteristic of biological systems. Its presence is necessary for life – as it underwrites an existential separation of the system from its environment, while preserving its interactions. The hierarchical organisation of complex systems – like living organisms – implies that the self-similar organisation of Markov blankets may be evident at any level of biological structure. From the point of view of dynamical systems, Markov blankets are attractors, attracting fast microscopic dynamics, while enforcing the emergence of macroscopic (order) parameters. This circular causality nicely captures the self-organisation of biological systems, which evolve autonomously from the environment in a morphology (Markov blanket) that is necessarily predisposed to a selective coupling with external states. The natural place – where these attractors might be specified – is the genetic code. Clearly, this is rather speculative; however, it is possible that the astonishing diversity in the flora and fauna we witness might reflect the fact that, in a world where signals are spatially dependent, Markov blankets are synonymous with existence.

### Methods

The simulations reported in this paper can be reproduced using the open access academic software SPM (<u>http://www.fil.ion.ucl.ac.uk/spm/software/</u>). The key routines are **DEM_cell.m** and **DEM_cell_cell.m** that illustrate self-organisation of a single ensemble and ensemble of ensembles respectively.

**DEM_cell.m:** This demo illustrates self-organisation in an ensemble of (sixteen) cells using the same principles described in **DEM_morphogenesis**, using a simpler generative model. Overall, the dynamics of these simulations show how one can prescribe a point attractor for each constituent of an ensemble that endows the ensemble with a point attractor to which the ensemble converges. In this example, we consider the special case where the point attractor is itself a Markov blanket. In other words, cells come to acquire dependencies, in terms of intracellular signalling, that conform to a simple Markov blanket with intrinsic or internal cells, surrounded by active cells that are, in turn, surrounded by sensory cells. This organisation rests upon intracellular signals and active inference using generalised (second-order) variational filtering. In brief, the hidden causes driving action (migration and signalling) are expectations about cell type. These expectations are optimised using sensory signals; namely, the signals generated by other cells. By equipping each cell with prior beliefs about what it would sense if it was a particular cell type (i.e., internal, active or sensory), the act (i.e., move and signal) so that they behave and infer their role in an ensemble of cells that itself has a Markov blanket. In a **DEM_cell_cell.m**, we use this first-order scheme to simulate hierarchical emergence of Markov blankets; i.e., ensembles of cells that can be one of three types at the local level; independently of their time at the global level.

**DEM_cell_cell.m:** This demo is a hierarchical extension of **DEM_cell.m**, where we have 16 ensembles comprising 16 cells. Each cell has a generative model (i.e., prior beliefs) about its local and global cell type (i.e., internal, active or sensory). Given posterior beliefs about what sort of self it is at the local and global level, it can then predict the local and global intracellular signals it would expect to receive. The ensemble of ensembles then converges to a point attractor; where the ensemble has a Markov blanket and each element of the ensemble comprises a cell that is itself a Markov blanket. The focus of this simulation is how the local level couples to the global level and vice versa. For simplicity (and computational expediency) we only model one ensemble at the local level and assume that the remaining ensembles conform to the same (local) dynamics. This is effectively a mean field approximation where expectations of a cell in the first ensemble about its global type are coupled to the corresponding expectations and the ensemble level, and vice versa. The results of this simulation are provided in the form of a movie and graphs.

## Additional information

### Author contribution

Ensor Rafael Palacios and Karl Friston conceptualized the idea, wrote the manuscript text and contributed to the simulations. Adeel Razi conceptualized the idea, reviewed the manuscript text and contributed to the simulations. Thomas Parr and Michael Kirchhoff reviewed the manuscript.

## References

[1] F. G. Varela, H. R. Maturana, and R. Uribe, “Autopoiesis: The organization of living systems, its characterization and a model,” BioSystems, vol. 5, no. 4, pp. 187–196, 1974.

[2] H. R. Maturana, “The Organization of the Living: A Theory of the Living Organization,” Int. J. Human-Computer Stud., vol. 51, no. June 1974, pp. 149–168, 1974.

[3] G. M. Whitesides, “Self-Assembly at All Scales,” Science (80-.)., vol. 295, no. 5564, pp. 2418–2421, 2002.

[4] G. Auletta, “A Paradigm Shift in Biology?,” Information, vol. 1, no. 1, pp. 28–59, 2010.

[5] K. Friston, “Life as we know it.,” J. R. Soc. Interface, vol. 10, no. 86, p. 20130475, 2013.

[6] S. A. Kauffman, The origins of order. 1993.

[7] S. Kauffman, At Home in the Universe: The Search for the Laws of Self-Organization and Complexity. 1995.

[8] G. F. R. Ellis, D. Noble, and T. O’Connor, “Top-down causation: an integrating theme within and across the sciences? Introduction,” Interface Focus, vol. 2, no. 1, pp. 1–3, 2012.

[9] E. Schrödinger, “What is Life? The Pysical Aspect of the Living Cell,” 1944.

[10] W. Tschacher and H. Haken, “Intentionality in non-equilibrium systems? The functional aspects of self-organized pattern formation,” New Ideas Psychol., vol. 25, no. 1, pp. 1–15, 2007.

[11] G. Nicolis and I. Prigogine, “Self-organization in nonequilibrium systems: from dissipative structures to order through fluctuations,” John Wiley Sons, p. 491, 1977.

[12] D. J. Evans and D. J. Searles, “The fluctuation theorem,” Adv. Phys., vol. 51, no. 7, pp. 1529–1585, 2002.

[13] K. Friston, K. Stephan, B. Li, and J. Daunizeau, “Generalised filtering,” Math. Probl. Eng., vol. 2010, 2010.

[14] K. Friston, “The free-energy principle: a unified brain theory?,” Nat. Rev. Neurosci., vol. 11, no. 2, pp. 127–138, 2010.

[15] R. C. Conant and W. R. Ashby, “Every good regulator of a system must be a good model of that system,” Int. J. Syst. Sci., vol. 1, no. 2, pp. 89–97, 1970.

[16] J. Hohwy, “The Self-Evidencing Brain.Noûs, 2016. 50(2):.,” Noûs, vol. 50(2), pp. 259–285, 2016.

[17] K. J. Friston, J. Daunizeau, J. Kilner, and S. J. Kiebel, “Action and behavior: A free-energy formulation,” Biol. Cybern., vol. 102, no. 3, pp. 227–260, 2010.

[18] A. Clark, Supersizing the Mind: Embodiment, Action, and Cognitive Extension. 2009.

[19] N. Ay, N. Bertschinger, R. Der, F. Güttler, and E. Olbrich, “Predictive information and explorative behavior of autonomous robots,” Eur. Phys. J. B, vol. 63, no. 3, pp. 329–339, 2008.

[20] J. Fuster, “Upper processing stages of the perception-action cycle,” Trends Cogn. Sci., vol. 8, no. 4, pp. 143–145, 2004.

[21] K. Friston, M. Levin, B. Sengupta, and G. Pezzulo, “Knowing one’s place: a free-energy approach to pattern regulation.,” J. R. Soc. Interface, vol. 12, no. 105, p. 20141383-, 2015.

[22] J. Pearl, “Probabilistic Reasoning in Intelligent Systems,” Morgan Kauffmann San Mateo, vol. 88. p. 552, 1988.

[23] G. Auletta, “Information and metabolism in bacterial chemotaxis,” Entropy, vol. 15, no. 1, pp. 311–326, 2013.

[24] K. Friston, B. Sengupta, and G. Auletta, “Cognitive dynamics: From attractors to active inference,” Proc. IEEE, vol. 102, no. 4, pp. 427–445, 2014.

[25] H. Crauel and F. Flandoli, “Attractors for random dynamical systems,” Probab. Theory Relat. Fields, vol. 100, no. 3, pp. 365–393, 1994.

[26] H. Crauel, S. Then, and S. Mathematik, “Global Random Attractors are Uniquely Determined by Attracting Deterministic Compact Sets,” Ann. di Mat. Pura ed Appl., vol. CLXXVI, no. Iv, pp. 57–72, 1999.

[27] K. Friston and P. Ao, “Free energy, value, and attractors,” Comput. Math. Methods Med., vol. 2012, 2012.

[28] T. D. Frank, “Nonlinear Fokker–Planck equations: fundamentals and applications”. Springer Series in Synergetics. Berlin, Germany: Springer, 2004.

[29] U. Seifert, “Stochastic thermodynamics, fluctuation theorems and molecular machines.,” Rep. Prog. Phys., vol. 75, no. 12, p. 126001, 2012.

[30] D. J. C. MacKay, Information Theory, Inference and Learning Algorithms, vol. 53, no. 9. 2003.

[31] H. Haken, Synergetics: An introduction. 1978.

[32] N. Balaskas et al., “Gene regulatory logic for reading the sonic hedgehog signaling gradient in the vertebrate neural tube,” Cell, vol. 148, no. 1–2, pp. 273–284, 2012.

[33] H. Y. Chang et al., “Diversity, topographic differentiation, and positional memory in human fibroblasts,” Proc. Natl. Acad. Sci., vol. 99, no. 20, pp. 12877–12882, 2002.

[34] A. Ochoa-Espinosa, D. Yu, A. Tsirigos, P. Struffi, and S. Small, “Anterior-posterior positional information in the absence of a strong Bicoid gradient,” Proc. Natl. Acad. Sci., vol. 106, no. 10, pp. 3823–3828, 2009.

